# Identification of the FSH-RH, the other gonadotropin-releasing hormone

**DOI:** 10.1101/2023.05.26.542428

**Authors:** Shun Kenny Uehara, Yuji Nishiike, Kazuki Maeda, Tomomi Karigo, Shigehiro Kuraku, Kataaki Okubo, Shinji Kanda

**Author notes:** These authors contributed equally to this work (Shun Kenny Uehara, Yuji Nishiike). Department of Genomics and Evolutionary Biology, Molecular Life History Laboratory, National Institute of Genetics, Shizuoka 411-8540, Japan (Shigehiro Kuraku).

## Abstract

In vertebrates, folliculogenesis and ovulation are regulated by two distinct pituitary gonadotropins: follicle-stimulating hormone (FSH) and luteinizing hormone (LH). Today, there is an intriguing consensus that a single hypothalamic neurohormone, gonadotropin-releasing hormone (GnRH), regulates the secretion of both FSH and LH, although the required timing and functions of FSH and LH are different. However, recent studies in vertebrates other than mammals indicate that the effect of GnRH on FSH is too weak to explain its regulation. Therefore, to challenge this “solo GnRH model,” we aimed to identify the other gonadotropin regulator, FSH-releasing hormone (FSH-RH), in vertebrates. Here, by using the model teleost medaka, we successfully identified cholecystokinin as the FSH-RH. Our histological and *in vitro* analyses demonstrated that hypothalamic cholecystokinin-expressing neurons directly affect FSH cells through the cholecystokinin receptor, Cckbr1, thereby increasing the expression and release of FSH. Remarkably, the knockout of cholecystokinin ligand or *cckbr1* minimized FSH expression and resulted in a complete failure of folliculogenesis. Our results challenge the longstanding consensus of the solo GnRH model in all vertebrates; instead, we propose the existence of a “dual GnRH model” group in vertebrates that utilizes both FSH-RH and LH-RH. The discovery of the FSH-RH in vertebrates opens not only a new era in neuroendocrinology but also possible applications involving vertebrate reproduction.

## Main Text

The pituitary gland secretes two essential gonadotropin hormones for gonadal functions: follicle-stimulating hormone (FSH) and luteinizing hormone (LH). Due to the importance in sexual development and reproduction in all vertebrates studied, the regulatory mechanism of these hormones has been extensively investigated. Importantly, given the hypothesis that the secretion of anterior pituitary hormones is mostly regulated by the hypothalamic neuropeptides ^1,2^, LH-releasing hormone (LH-RH) was discovered in 1971 as the key regulator of LH and FSH release in mammals, marking a significant breakthrough ^1,3,4^. LH-RH was later studied in other mammals, and its involvement in reproductive regulation has been generally established in vertebrates ^5^. Consequently, it has been widely recognized that LH-RH is the sole gonadotropin-releasing hormone (GnRH), responsible for regulating both FSH and LH secretion in vertebrates ^6^, and it is now referred as the final common pathway for central regulation of gonadotropin secretion or fertility ^7-9^.

However, this once-established consensus has been challenged in vertebrates other than mammals. Intriguingly, it has been reported that GnRH knockout (KO) does not affect FSH function in model teleosts such as medaka and zebrafish ^10,11^, which implies that GnRH may not be the primary regulator of FSH release, at least in teleosts. Also, in quail, a model avian species, reports clearly state that GnRH effects on FSH are smaller than those on LH release ^12,13^. This observation raises the same perplexing mystery indicated in teleosts: that GnRH has only a small effect on FSH as observed ^1,14-16^. Thus, it is postulated that an exclusive FSH-regulating hormone, FSH-RH, exists in vertebrates alongside GnRH. This hypothesis suggests that a significant number of vertebrates, rather than being limited to minor subgroups, may utilize distinct FSH-RH and LH-RH systems, indicating the presence of a “dual GnRH model.”

In the present study, to challenge the current “solo GnRH model,” we aimed to identify the FSH-RH. Among non-mammalian vertebrates, we used medaka because of their amenability to genetic modification and the exceptionally accessible information about their reproductive biology ^9,17-20^. The use of medaka is also advantageous because, as teleosts, they have separate cells that express FSH or LH, unlike mammals, in which FSH and LH are secreted from a single cell type, making medaka useful for examination of the FSH-RH receptors ^21-25^.

### Cckbr1 is highly expressed in FSH cells and essential for FSH function

First, considering the importance of identifying the receptor expressed in the FSH cells, we performed FSH-cell specific RNA-sequencing (RNA-seq) by collecting green fluorescent protein (GFP)-positive cells in the pituitary of FSH-GFP medaka (*n* = 4; Extended Data Fig. 1). Among the metabolic receptors expressed, we found that cholecystokinin B receptor 1 (*cckbr1*) had the highest expression (Extended Data Table 1). Therefore, we conducted double *in situ* hybridization of *FSH subunit beta gene, fshb*, and *cckbr1* to establish their co-expression in the pituitary through the histological method. From these experiments, we demonstrated that *cckbr1* is expressed in FSH cells (Fig. 1a). To determine the functional essentiality of the identified receptor, we generated *cckbr1* KO and analyzed phenotypes in both females and males. Our analysis revealed that both *cckbr1*^−/−^ females and males have substantially smaller gonads compared to *cckbr1*^+/+^ or *cckbr1*^+/−^ individuals (Fig. 1b). In addition, a severe reduction in gonadal weight (normalized by body weight, gonadosomatic index), was observed in both *cckbr1*^−/−^ females and males (*n* = 5; Fig. 1 c,d), whereas there was no apparent change in body weight (Extended Data Fig. 2). These results suggest that the same mechanism is established regardless of sex. Histological analysis revealed that the ovaries of the *cckbr1*^−/−^ females include only the previtellogenic oocyte and they are infertile, whereas wild-type ovaries show full-grown oocytes, indicating fertility (Fig. 1e). Unlike females, *cckbr1*^−/−^ males showed normal spermatogenesis (Fig. 1e), which is similar to the results of *fshb* KO medaka, in that only females show infertility ^10^. Estradiol (E2) content of the whole body of the female (*n* = 3) was analyzed in *cckbr1*^*+/+*^ and *cckbr1*^−/−^ fish and indicated that the E2 concentration was significantly lower in *cckbr1*^−/−^ than in *cckbr1*^*+/+*^ fish (Fig. 1f). Finally, the pituitaries of *cckbr1*^*+/*−^ and *cckbr1*^−/−^ fish (*n* = 7) were subjected to quantitative reverse transcription polymerase chain reaction (qRT-PCR) to examine the effects of KO on gonadotropin gene expression in females and males (Fig. 1g-l). In both sexes, *fshb* expression was much lower in *cckbr1*^−/−^ than in *cckbr1*^*+/*−^ (Figure 1g,j). Interestingly, only the expression of *LH subunit beta* gene (*lhb*) in females was lower in *cckbr1*^−/−^ than in *cckbr1*^+/−^ (Fig. 1h,k). To diminish the secondary effect from the immature gonad due to the reduction in FSH secretion, we compared the expression of KOs after ovariectomy. In the ovariectomized females (*n* = 6), there was no difference in *lhb* expression between *cckbr1*^*+/*−^ and *cckbr1*^−/−^ (Extended Data Fig. 3). These results strongly suggest that the difference in *lhb* expression between intact *cckbr1*^*+/*−^ and *cckbr1*^−/−^ females should be due to the secondary effect of reduced serum estrogen concentration in *cckbr1*^−/−^ because ovarian estrogen strongly enhances the expression of *lhb* ^24^. Expression of *thyroid stimulating hormone subunit beta* gene (*tshb*) did not change in either sex (Fig. 1i,l). Thus, we proved that Cckbr1, which is expressed in FSH cells, is essential for the normal function of FSH ^1,24^.

**Fig. 1.**
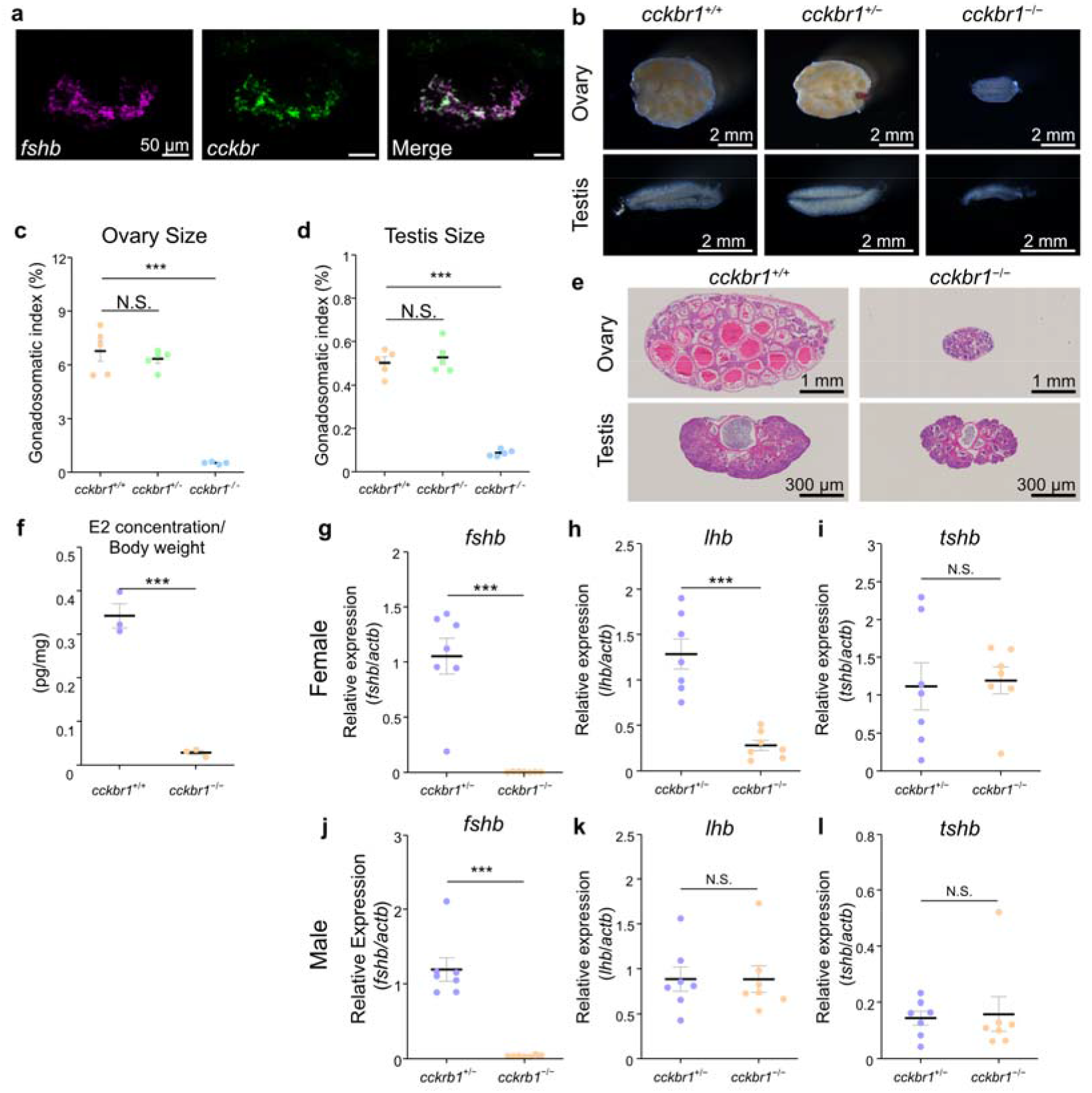
Cckbr1 expressed in the pituitary FSH cells is essential for normal FSH expression. **a**, Double *in situ* hybridization of *fsh* (magenta) and *cckbr1* (green) in the pituitary of medaka indicating their co-expression. **b-d**, Knockout of *cckbr1* showing abnormally smaller gonads in both females and males. Gross morphology of gonads (b) as well as ovarian size (c) and testis size (d) show a drastic reduction in *cckbr1*^*−/−*^ in both sexes. **e**, Histological sections indicate the severe effect of knockout in the ovary and testis. **f**, The estradiol (E2) content of females shows a drastic reduction in *cckbr1*^−/−^ compared to the wild-type medaka. **g-l**, Expression of *fshb* (g, j), *lhb* (h, k), and *tshb* (i, l) in the pituitary of each genotype analyzed by qRT-PCR. Drastically reduced *fshb* mRNA was observed in *cckbr1*^−/−^ in both females (g) and males (j). *lhb* mRNA showing a reduction in *cckbr1*^−/−^, which may be due to the reduced serum estradiol in females (h), whereas showing no effect of KO in males (k). *tshb* mRNA between *cckbr1*^*+/*−^ and *cckbr1*^−/−^ in females (i) and males (l). \*\*\**P*<0.001, N.S. not significant.

### Ligand for Cckbr1 (cholecystokinin) is expressed in the hypothalamic neurons that project to the pituitary

After identifying Cckbr1 as the prime candidate for the FSH-RH receptor, the ligands for Cckbr1 were explored. In vertebrates, genes for ligands that bind to Cckbr1 are members of the gastrin/cholecystokinin family; medaka have two cholecystokinin (CCK) paralogs: *cholecystokinin a* (*ccka*) and *cholecystokinin b* (*cckb*), as well as *gastrin* ^26^. Because the highly conserved 8-amino acid peptide of CCK and Gastrin are reported to show high biological activity, we used CCK8 (DY(SO_3_H)LGWMDF-_NH2_) and Gastrin8 (DY(SO_3_H)RGWLDF-_NH2_) for the luciferase reporter assay. Note that both *ccka* and *cckb* genes result in the identical deduced 8-amino acid residue peptide CCK8. In the luciferase assay, for all cyclic adenosine monophosphate (cAMP), Ca^2+^, and mitogen-activated protein kinase (MAPK) reporter systems, Cckbr1 was activated by both CCK8 and Gastrin8 in dose-dependent manners (Fig. 2a). The half-maximal effective concentrations (EC_50_) are summarized in Extended Data Table 2. Thus, CCK and Gastrin can activate the receptor, Cckbr1, at physiological concentrations.

**Fig. 2.**
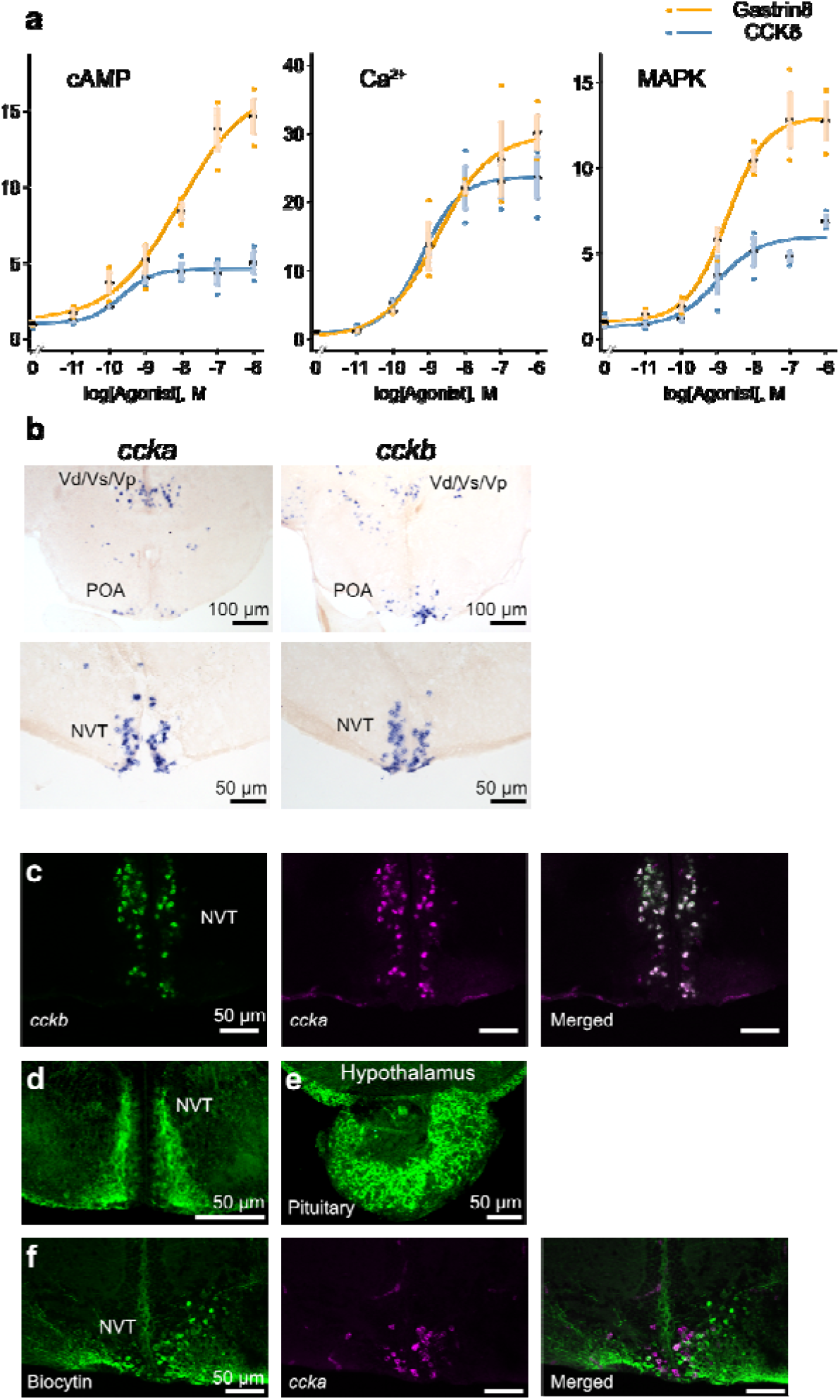
Hypophysiotropic neurons expressing *ccka* and *cckb* exist in the hypothalamus. **a**, Luciferase assay of cholecystokinin family peptides CCK8 and Gastrin8 for cAMP, Ca^2+^, and MAPK pathways in HeLa cells expressing Cckbr1. **b**, *in situ* hybridization of *ccka* and *cckb* in the pre-optic area (POA) and the nucleus ventralis tuberis (NVT) of the hypothalamus. **c**, Double *in situ* hybridization of *ccka* (magenta) and *cckb* (green) in the NVT. **d**–**e**, Immunohistochemistry (IHC), using a CCK-specific antibody labeling CCK neurons in the NVT (d) and axonal fibers in the pituitary (e). **f**, Dual labeling of retrogradely induced biocytin (green) from the pituitary and *ccka* mRNA (magenta).

The ligands were further analyzed for their expression in the hypothalamus. Because reverse transcription PCR (RT-PCR) of the whole brain indicated high expression of *ccka* and *cckb* but not *gastrin* (Extended Data Fig. 4), we examined the localization of their mRNAs in the brain by *in situ* hybridization, using *ccka* and *cckb* probes to analyze their possible expression. Here, *ccka* and *cckb* were expressed in various regions of the brain including the nuclei that contain the hypophysiotropic (pituitary-projecting) neurons, the pre-optic area (POA), and the nucleus ventralis tuberis (NVT) in the hypothalamus (Fig. 2b). Because the expression of *ccka* and *cckb* in the NVT apparently overlapped, we examined the possible co-expression of *ccka* and *cckb* by double *in situ* hybridization. We discovered that the vast majority of CCK neurons in NVT co-express *ccka* and *cckb* mRNA (Fig. 2c). To observe their projection, we used a commercial cholecystokinin antibody. As expected from previous studies of comparative anatomy ^27,28^, cell bodies in the nuclei including the hypothalamus (Fig. 2d), as well as dense axonal fibers in the pituitary (Fig. 2e), were labeled. Note that all cell bodies and fibers were diminished in the double knockout of *ccka* and *cckb*; the cell bodies and axonal projection in the pituitary are strongly suggested to originate from *ccka*- and (or) *cckb*-expressing neurons (Extended Data Fig. 5). Next, to examine which population of these neurons projected to the pituitary, we performed double labeling of *ccka* by *in situ* hybridization and retrograde tracer labeling from the pituitary. In the sample that underwent biocytin tracer labeling in the area where FSH cells are localized in the pituitary, we observed co-labeling of retrogradely labeled biocytin and *ccka* mRNA in the neurons in the NVT (Fig. 2f). These results strongly suggest that the CCK neurons in the NVT are hypophysiotropic and project their axons to the pituitary FSH cells.

### *in vitro* experiments suggest that CCK increases both the release and expression of FSH

To examine the possibility that the release of FSH is induced by CCK, Ca^2+^ imaging of FSH cells was performed. The pituitaries of females were isolated from the *fsh*:inverse pericam (*fsh*:IP) transgenic medaka ^29^, whose FSH cells express a genetically encoded Ca^2+^ indicator called inverse pericam ^30^. In FSH cells, 1 μM CCK8 induced a rapid and strong intracellular Ca^2+^ increase, which triggers hormonal release from the FSH cells (Fig. 3a). This effect was also observed in males (Extended Data Fig. 6). Perfusion of various concentrations of CCK8 (*n* = 4) demonstrated that this action on FSH cells was dose-dependent (EC_50_ = 19 nM, Fig. 3b). Unlike that in FSH cells, the Ca^2+^ imaging experiment in LH cells using the *lh*:IP transgenic medaka ^29^ (*n* = 4) did not show a CCK8-induced Ca^2+^ response (Fig. 3c,d). We concluded that CCK acts upon FSH cells, but not on LH cells, to induce hormonal release.

**Fig. 3.**
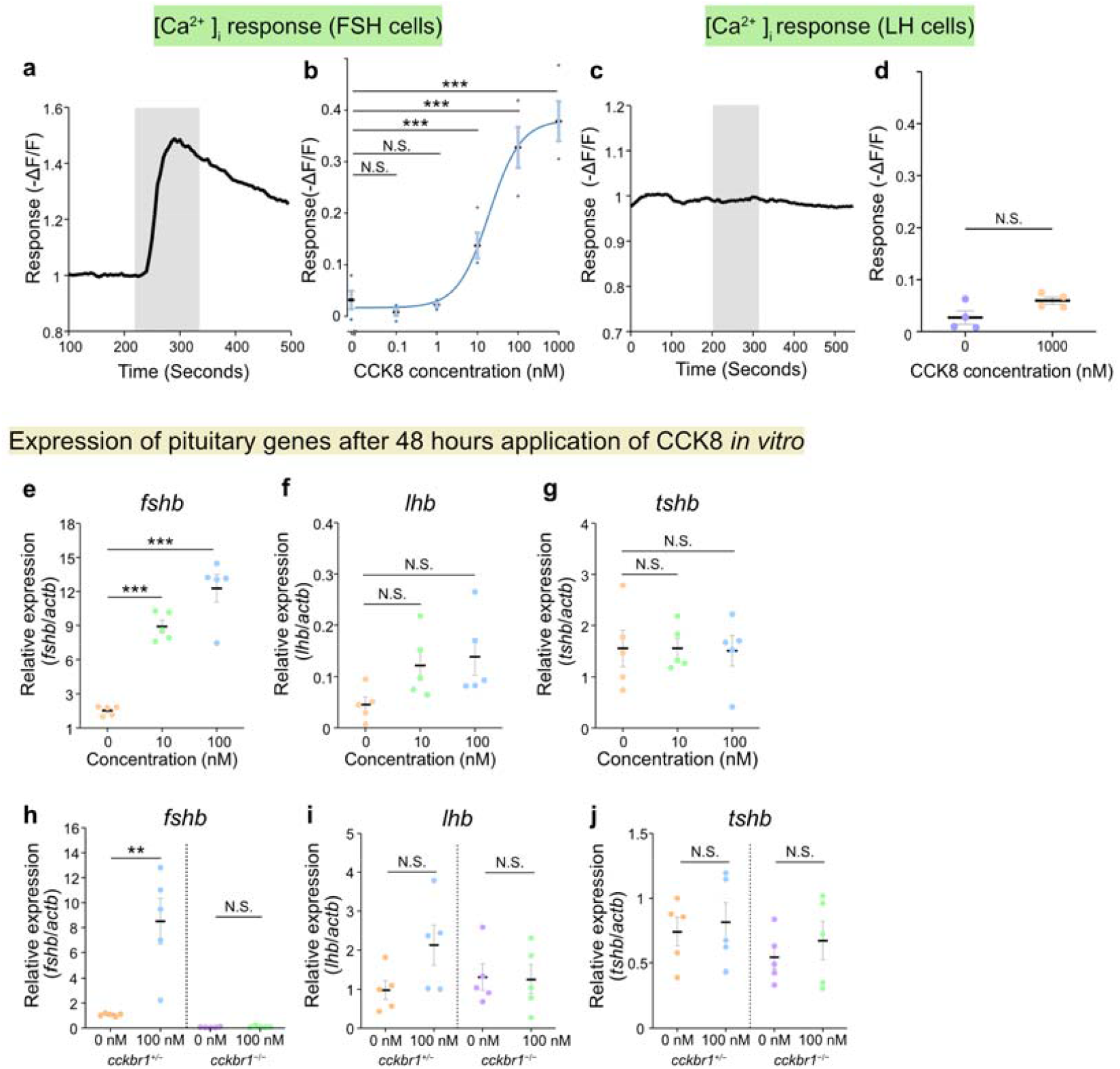
CCK robustly increased both [Ca^2+^]_i_, which causes the hormonal release, and mRNA expression of *fshb*. **a**, Ca^2+^ imaging of FSH cells applied with 1 μM CCK8. **b**, Dose-response curve of [Ca^2+^]_i_ in FSH cells when applied with 0, 0.1, 1, 10, 100, or 1000 nM CCK8. **c**, Ca^2+^ imaging of LH cells applied with 1 μM CCK8. **d**, Repetitive Ca^2+^ imaging trials on LH cells. **e**–**g**, qRT-PCR of the pituitary after *in vitro* incubation with CCK8 for 48 hours. \*\*\**P*<0.001, N.S., not significant. **h**–**j**, The effects of CCK8 using *cckbr1*^*+/*−^ and *cckbr1*^−/−^ pituitary (h, *fshb*; i, *lhb*; j, *tshb*). \*\**P*<0.01.

We next examined if CCK stimulates FSH expression by an *in vitro* experiment of isolated pituitary. The pituitaries of female medaka (*n* = 5) were isolated and incubated for 48 hours in a culture medium containing 0, 10, or 100 nM CCK8. After incubation, the expression of *fshb, lhb*, and *tshb* mRNA was analyzed by qRT-PCR, which revealed that *fshb* expression was dose-dependently increased (Fig. 3e-g). In the presence of 10 or 100 nM CCK8 during incubation, *fshb* increased approximately 10-fold compared to the 0 nM control (Fig. 3e). Expression of *lhb* mRNA showed a slight but nonsignificant increase (Fig. 3f). Considering the results of Ca^2+^ imaging and double *in situ* hybridization, LH cells likely do not possess CCK receptors. Thus, this may be an indirect effect. Also, CCK8 did not affect *tshb* expression (Fig. 3g). A similar procedure was conducted using the pituitaries of *cckbr1*^*+/*−^ and *cckbr1*^−/−^ medaka. As expected, CCK8 increased *fshb* expression in the *cckbr1*^*+/*−^ pituitary but not in the *cckbr1*^−/−^ pituitary (Fig. 3h). These results support that the action of CCK on FSH cells is mediated by Cckbr1. Similar to the results in the wild-type medaka, CCK8 did not significantly increase the expression of *lhb* or *tshb* in both *cckbr1*^*+/*−^ and *cckbr1*^−/−^ genotypes (Fig. 3i,j).

### CCK is essential for FSH function

From the results above, we showed that CCK strongly stimulated FSH secretion, both hormone synthesis and release. To examine the essentiality of intrinsic CCK, we generated KOs of the two CCK paralogs, *ccka* and *cckb*, by CRISPR ^31,32^. Interestingly, although a single KO of *ccka* or *cckb* resulted in a normal phenotype, the double KO showed a severe change in phenotype (Fig. 4, Extended Data Fig. 7). The overall phenotype of the double KO was similar to that of the *cckbr1* KO. In both female and male, the gonadal size of *ccka/cckb* double KO was drastically decreased (Fig. 4a,b, Extended Data Fig. 7). As expected, pituitary mRNA expression (*n* = 6) of *fshb* in *ccka/cckb* double KO was much smaller than that of the other genotypes (Fig. 4c). Also, *lhb* expression decreased in the double KO, whereas *tshb* did not differ among all genotypes (Fig. 4d,e). The fact that only the double KO was associated with a severe phenotype suggests that *ccka* and *cckb* have redundancy in their function of FSH regulation, which is also consistent with the fact that *ccka* and *cckb* co-localize in the NVT hypophysiotropic neurons (Fig. 2).

**Fig. 4.**
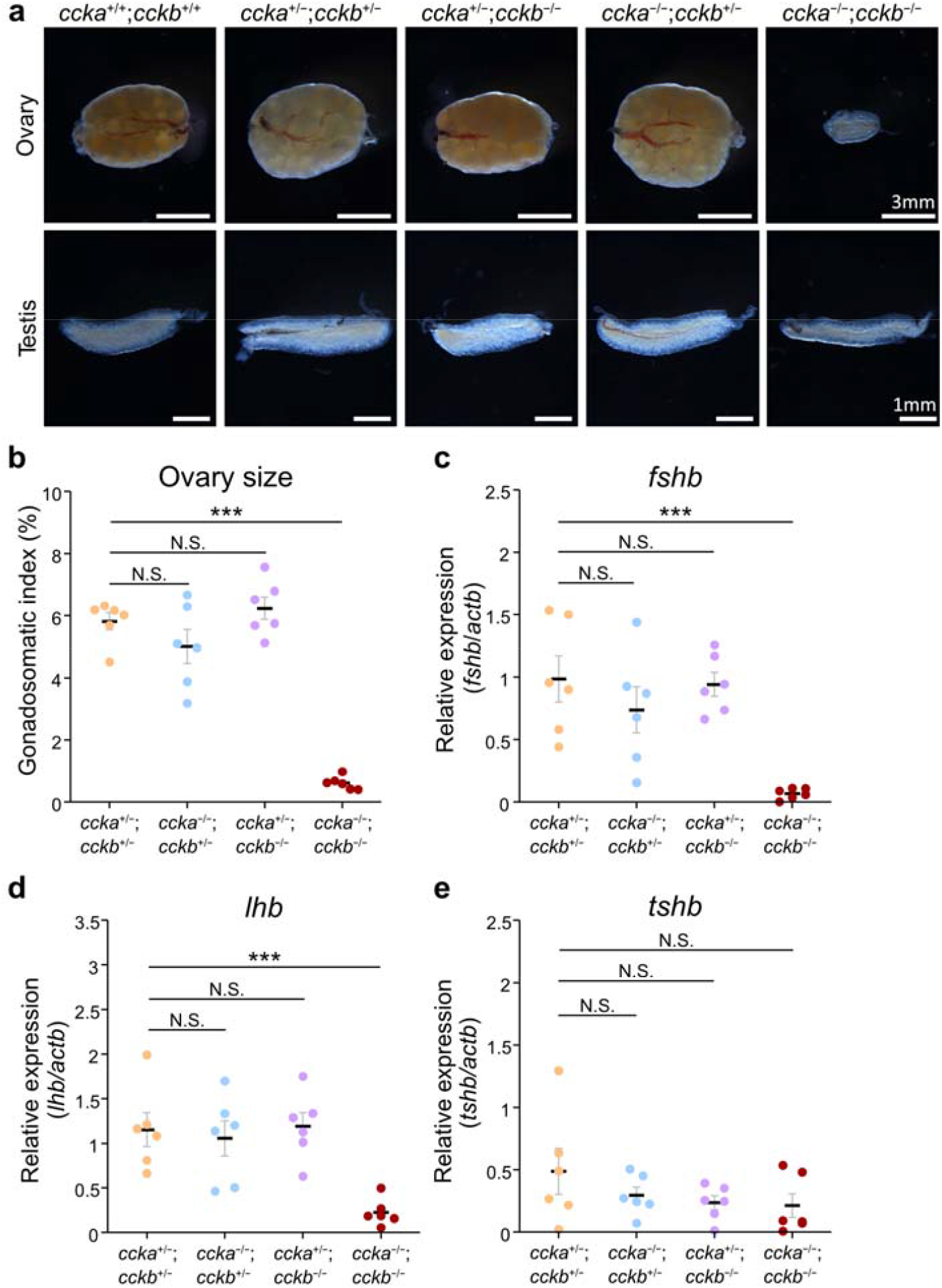
CCK is essential for FSH function. **a**, Gross morphology of the ovaries and testes of single and double KO of *ccka* and *cckb*. **b**, The ovarian size of each genotype normalized by body weight. **c-e**, qRT-PCR for *fshb, lhb*, and *tshb* of each genotype in females. \*\*\**P*<0.001, N.S., not significant.

## Discussion

In contrast to the current solo GnRH model hypothesis established in mammals, we identified hypothalamic CCK as the most potent and essential regulator of FSH release. The strong effects shown in *in vitro* experiments and the severe phenotype in knockout experiments strongly suggest that CCK is the FSH-RH. It was surprising that the long-absent FSH-RH was proven to be CCK, which is well known as an intestinal peptide whose function has been extensively studied in the mammalian digestive system in both basic and clinical research contexts ^33^. To date, despite many trials mainly in rodent models, a strong regulator of FSH other than GnRH has not been identified, thus it was hypothesized that LH-RH/GnRH is the only gonadotropin-releasing hormone in vertebrates. However, our discovery of FSH-RH provides strong counterevidence of this hypothesis. The evidence provided in this study suggests the existence of species in which FSH and LH are regulated by distinct hypothalamic factors, namely the “dual GnRH model” (Fig. 5a).

**Fig. 5.**
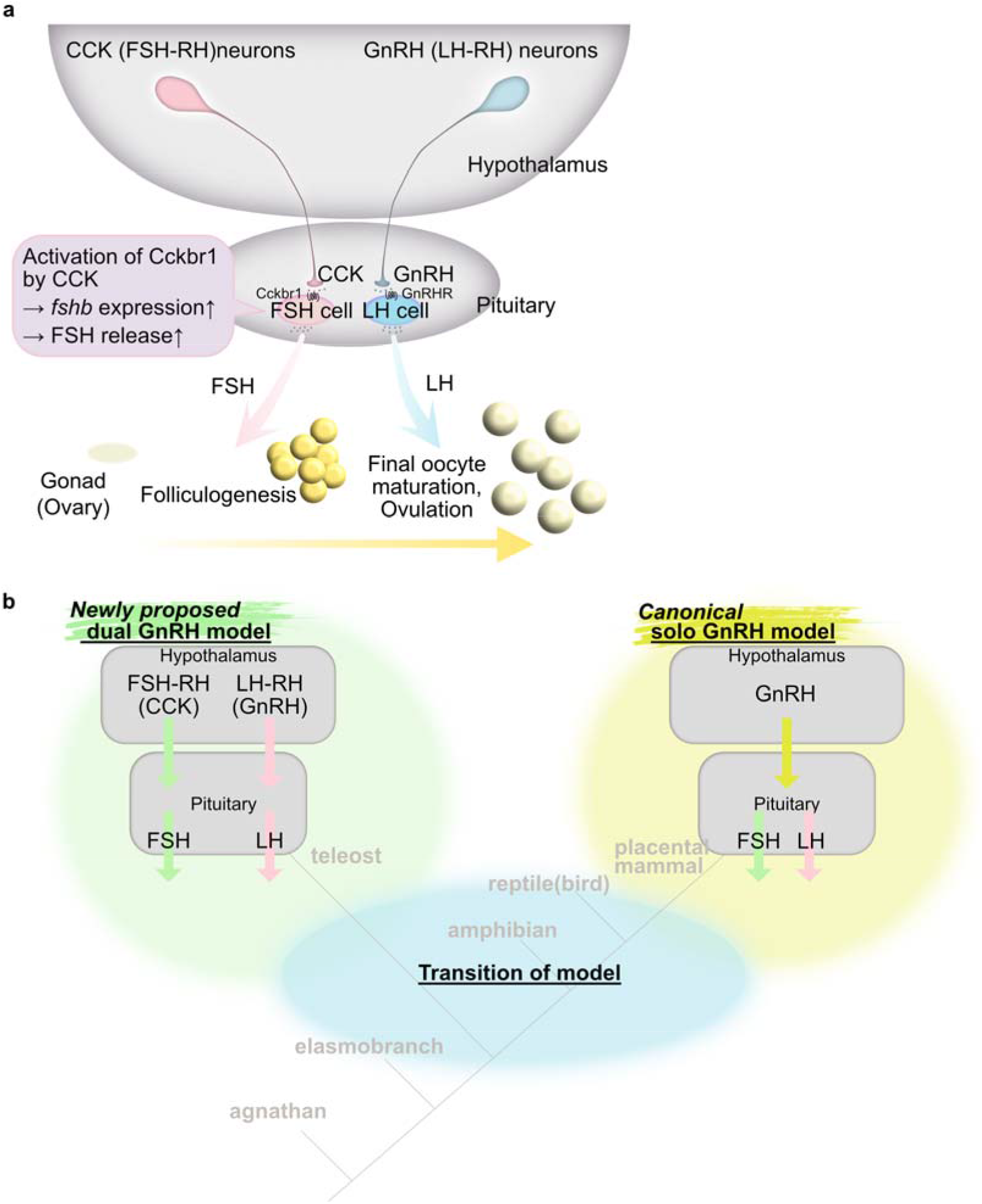
Working hypothesis of the “dual GnRH model” in which two gonadotropins are regulated by two gonadotropin-releasing hormones, FSH-RH and LH-RH. **a**, The present study identified CCK as the hypothalamic FSH-RH, which stimulates the expression and release of FSH to induce folliculogenesis. After folliculogenesis is completed, GnRH (LH-RH)-induced LH induces final oocyte maturation followed by ovulation. **b**, We propose a “dual GnRH model” group in vertebrates, in addition to the canonical solo GnRH model group. Phylogenetic relationships indicate that a transition of the models occurred during vertebrate evolution.

Although this study provides evidence suggesting that CCK acts as the FSH-RH in medaka, there are several indications that this mechanism could extend to other vertebrates. First, its mechanisms are likely to be conserved in all teleosts, at least. A cell type-specific RNA-seq study indicated that FSH-expressing cells showed high expression of CCK receptor in another teleost, tilapia ^34^. Moreover, in the Japanese eel, which is known to have diverged from other teleosts at the earliest stages of the teleost lineage ^35^, we demonstrated that *cckbr* mRNA is expressed in FSH cells (Extended Data Fig. 8). Therefore, the expression of CCK receptors in FSH cells appears to be widely conserved, at least in teleosts. Furthermore, regarding the CCK ligand, some immunohistochemical studies have suggested the existence of CCK immunoreactive neurons that project to the pituitary in the hypothalamus of other teleosts ^27,28^. Additionally, the fact that hypophysiotropic CCK neurons co-expressed the paralogous *ccka* and *cckb* in medaka, as shown in the present study, strongly suggests that the enhancer that enabled the expression of CCK in the hypophysiotropic neurons was acquired before the divergence of *ccka* and *cckb* genes, which occurred in the teleost-specific whole genome duplication ^36^. This can be further interpreted as indicating that the ancestral vertebrates, before the emergence of teleosts, also possessed hypophysiotropic CCK neurons in their hypothalamus. These lines of evidence suggest that the mechanism proved in this study is widely conserved, at least in teleosts and even in other lineages. It is also worth noting that the CCK receptor is highly expressed in the chicken pituitary ^37^. These facts raise the possibility of broad conservation of the “dual GnRH model” across vertebrates.

On the other hand, in placental mammals, there is currently no evidence to refute the established belief that they follow the solo GnRH model. In rodents, CCK does not induce FSH secretion in rats ^38^, and CCK-deficient mice are fertile ^39^. Furthermore, the *Gnrh*-deficient mouse shows a reduction in serum FSH concentration ^40^, indicating that LH-RH is an essential regulator for FSH as well as LH in this group. However, it should be inherently complex for the solo regulator to differentially regulate the release of distinct hormones. Perhaps these multiple roles of GnRH could be accomplished through the action of kisspeptin neurons. These neurons facilitate multi-modal GnRH secretion patterns, specifically the pulsatile and surge modes ^41,42^. While it is generally understood that the regulation of GnRH neurons by kisspeptin is unique to mammals, as birds lack kisspeptin-related genes ^25,43^ and knockout teleosts can reproduce ^44,45^, the mechanism behind the differential secretion of FSH and LH in all non-mammalian vertebrates remains unexplained. Thus, the dual GnRH model could potentially elucidate the mechanism in species beyond just teleosts. Moreover, there exists an intriguing report indicating CCK-induced FSH release in human pituitary tumors ^46^, implying the possibility that regulation of FSH by CCK may have existed in our ancestors.

Identification of the FSH-RH in the present study paves the way for new research avenues in neuroendocrinology. Given the evidence in mammals and teleosts, there should be a transition between the “dual GnRH model” and “solo GnRH model” in vertebrate evolution (Fig. 5b). It is crucial to understand which model the common ancestor of mammals and teleosts possessed. Further examination of vertebrate species should reveal the origin and evolutionary trajectories of the dual GnRH model and the solo GnRH model in vertebrates.

## Materials and Methods

### Animals

Medaka (*Oryzias latipes*) were maintained and used in accordance with the guiding principle for the Use and Care of Experimental Animals of the University of Tokyo. Male and female medaka (d-rR strain) and all the genetically modified medaka were maintained under a 14-hour light and 10-hour dark photoperiod at a water temperature of 28 °C. The fish were fed at least twice a day with live brine shrimp and flake food except for Sunday. In each subsequent experiment, siblings raised under the same conditions were used to control for genetic and environmental factors. Japanese eels (*Anguilla japonica*) were purchased from commercial source. Experiments were conducted in accordance with the protocols approved by the animal care and use committee of the University of Tokyo (Permission number, P22-6, P22-7).

### FSH-cell Specific RNAseq to find the expressed receptors

To find highly expressed receptors in FSH cells, we performed FSH-cell specific RNAseq using GFP-labeled transgenic medaka. First, we established a transgenic medaka whose FSH cells are labeled by EGFP. After injecting a plasmid construct containing ∼2 kb of 5’ flanking region of *fshb* of medaka and medaka heat shock protein minimal promoter (0.8 kb), the transgenic founder was selected in the F1 generation.

By using this transgenic medaka, FSH cells were collected based on their GFP fluorescence (Extended Data Fig. 1). First, pituitaries were dissected and subjected to dissociation with 0.2 mg/ml collagenase in artificial cerebrospinal fluid (ACSF) for 30 mins. By gentle pipetting with a narrowed pipette tip, the pituitary cells were dissociated. Under an inverted fluorescent microscope (AxioVert 100, Zeiss) with a manipulator, GFP-positive cells were collected with a glass pipette. Up to ten GFP-positive cells were pooled to each biological replicate.

After collecting four samples, sequence samples for Illumina sequencers were prepared using NEBNext Single Cell/Low Input RNA Library Prep Kit for Illumina (New England Biolabs, Ipswich, MA) according to the manufacturer’s instruction. Sequencing of the resulting sequence samples was outsourced to a commercial sequencing service using Novaseq 6000 (Nippon Genetics, Tokyo, Japan). Sequence data were subjected to mapping and read count by STAR and R-SEM, respectively. The genes with receptors in the annotation name were selected and sorted in descending order.

### Single and double in situ hybridization

Adult female d-rR wildtype medaka were deeply anesthetized with 0.02% tricaine methanesulfonate (MS-222) and cryosections were prepared according to the protocol previously reported ^20^ at 25 μm with a cryostat (Leica CM3050, Leica, Wetzlar, Germany; objective temperature: -24°C, chamber temperature: -28°C). The sliced sections were placed on the coated slide glass (CREST-coated, Matsunami, Kishiwada, Japan).

We analyzed the distribution of *cckbr1, ccka*, and *cckb* messenger RNA (mRNA) by *in situ* hybridization (ISH). Also, double *in situ* hybridization was performed to examine the co-localization of two genes. Single and double *in situ* hybridization was conducted in the protocol as described previously ^44^. *cckbr1*-, *ccka*-, and *cckb*-specific digoxigenin (DIG)-labeled mRNA probes, and *cckb*- and *fshb*-specific Fluorescein-labeled mRNA probes were prepared based on the cDNA region amplified by the primers listed in Extended Data Table 3 (Gene names in Ensembl: *ccka*, ENSORLG00000005949; *cckb*, ENSORLG00000005594; *cckbr1*, ENSORLG00000017966). We also used a *fshb* probe as described previously ^24^. The signal of single ISH was visualized by NBT/BCIP. After the coverslip, we observed the sections through BX53 biological microscope (Olympus, Tokyo, Japan). It was confirmed that signals are observed in slides hybridized with antisense probes and are not observed in sense probes (Extended Data Fig. 9). We followed the Medaka Histological Atlas for the nomenclature of the medaka brain nuclei. Double *in situ* hybridization was visualized by TSA plus biotin followed by ABC Elite kit (Vector Laboratories, Burlingame, CA) and Streptavidin, Alexa Fluor 488 (Green; Thermo Fisher, Waltham, MA), and TSA plus Cy3 (Red; Akoya Bioscience, Marlborough, MA). Note that the first peroxidase label on the fluorescein probe was completely quenched with 3% H_2_O_2_ for 40 mins before the second antibody for the DIG probe was labeled. The same protocol was applied to Japanese eel single/double *in situ* hybridization with minor modifications. The body weight of Japanese eel were 260-290g. Probes were based on the database in the National Center for Biotechnology Information (NCBI). NBCI reference sequences are as follows. *eel_fshb*, XM_035417932.1; *eel_cckbr*, XM_035394241.1. The primers used to prepare probe templates are listed in Extended Data Table 3. We observed the signals through a confocal microscope FV-1000 (Olympus) or Leica TCS SP8 (Leica Microsystems).

### Immunohistochemistry

Adult female d-rR wildtype medaka were deeply anesthetized and their brains were fixed with 4% paraformaldehyde (PFA) in PBS and cryosectioned. Their body weight is 0.11-0.15g. To label the CCK-expressing cells, we used an anti-cholecystokinin (26-33) antibody raised in rabbit (C2581, Sigma-Aldrich, St. Louis, MO; 1:5,000). After antigen retrieval with HistoVT (Nacalai Tesque, Kyoto, Japan) according to the manufacturer’s protocol, a primary antibody was applied with 5% normal goat serum. After incubation with anti-rabbit IgG, a secondary antibody, signal amplification with an ABC Elite kit (Vector Laboratories, Burlingame, CA) was applied. Immunoreactivities were visualized with Alexa Fluor 488-conjugated streptavidin (1:500; Invitrogen). Some of the samples were labeled with 4’,6-diamidino-2-phenylindole (DAPI). We observed the signals through a confocal microscope FV-1000 (Olympus) or Leica TCS SP8 (Leica Microsystems).

### Dual labeling of Biocytin and ccka mRNA

Adult female *fsh*:Inverse Pericam transgenic medaka (*fsh*:IP) ^29^ were used in this experiment to visualize the location of FSH cells. They were deeply anesthetized with 0.02% MS-222. The fish was decapitated, and the brain was excised. The fluoresced area (around FSH cells) of the pituitary was carefully injected with biocytin using the glass needle prepared from a glass capillary (G-1.5; Narishige, Tokyo, Japan) with a micropipette puller (P-97; Sutter Instruments, Novato, CA). The brain was incubated with the artificial cerebral spinal fluid (ACSF) containing (in mM): 134 mM NaCl, 2.9 mM KCl, 1.2 mM MgCl_2_, 2.1 mM CaCl_2_, 10 mM HEPES, and 15 mM glucose (pH 7.4, adjusted with NaOH) for 30 minutes and fixated with 4% PFA in PBS, then 30% sucrose in PBS and cryosectioned as mentioned above. Biocytin signal was amplified and visualized using ABC kit and Streptavidin, Alexa Fluor 488-conjugated (1:500). *ccka* mRNA signal was detected by ISH using a DIG-labelled probe and visualized using anti-DIG POD antibody (1:500) and TSA plus Cy3.

### Generation of knockout medaka

*ccka, cckb*, and *cckbr1* knockout medaka were generated by CRISPR/Cas9-mediated genome editing. For *cckbr1* knockout, a CRISPR RNA (crRNA) was designed to target exon 1; for *ccka* knockout, two crRNA were designed to target exon 2, which encodes the signal peptide of the Ccka precursor protein, and exon 3, respectively; for *cckb* knockout, two crRNA were designed to target exon 3 and exon 4, which encodes the mature peptide of the CCK8, respectively. crRNA and trans-activating CRISPR RNA (tracrRNA) were synthesized by Fasmac (Kanagawa, Japan).

Target sequences of the CRIPSR RNA including PAM are as follows. *cckbr1*, AAGCGTGGACGGGTTCACGCAGG; *ccka*, TGACGCGTGTGATTGGTTAGTGG; *cckb*, GGAGTGCTGGCCCTCATCTGAGG. The crRNA, tracrRNA, and Cas9 protein (Nippon Gene Co. Ltd., Tokyo, Japan) were co-microinjected into medaka embryos at the one-cell or two-cell stage. Potential founder fish were screened by outcrossing with wildtype fish and testing progeny for mutations by direct sequencing. For the *cckbr1* knockout line, a founder was identified that produced progeny carrying an 8-bp deletion that caused frameshift leading to complete loss of transmembrane domains. For the *ccka* knockout line, a founder was identified that produced progeny carrying a 743-bp deletion that caused frameshift leading to premature truncation of the Ccka precursor protein. For the *cckb* knockout line, a founder was identified that produced progeny carrying a 210-bp deletion that caused frameshift leading to premature truncation of the Cckb precursor protein. These progenies were intercrossed to establish knockout lines. *ccka*/*cckb* double-knockout medaka were generated by crossing the progenies from the *ccka* knockout line and *cckb* knockout line. Each line was maintained by breeding heterozygous or double-heterozygous individuals to obtain wildtype, heterozygous, and knockout siblings for experimental use. The genotype of each fish was first determined by direct sequencing and thereafter by PCR and high-resolution melting analysis (for *cckbr1* knockouts) or sequencing, or agarose gel electrophoresis (for *ccka* and *cckb* knockouts) using the primers listed in Extended Data Table 3. The exon-intron structure and the deletion are illustrated in the Extended Data Fig. 10.

### E2 measurement

Peripheral tissues (the caudal halves of the bodies) were collected from *cckbr1*^+/+^and *cckbr1*^-/-^ females at 2–4.5 hours after the onset of the light period, frozen at -80°C, and homogenized with Micro Smash (Tomy Seiko Co. Ltd., Tokyo, Japan). Tissue lipids were extracted with diethyl ether. Tissue levels of E2 were determined using the Estradiol ELISA Kit (Cayman Chemical Company, Ann Arbor, MI). Peripheral levels of E2 were expressed as picograms per mg tissue weight.

### Gonadal size and histology

After recording the body weight of each of the males/females (from the *cckbr1* knockout line or *ccka*/*cckb* double-knockout line), the gonad was removed, weighed, and photographed under a stereo microscope M205FA (Leica Microsystems) equipped with a digital camera DFC7000T (Leica Microsystems). The gonadosomatic index (GSI) was calculated by the following formula. GSI = [gonad weight / total tissue weight] × 100. After photographed, the gonads were fixed in 4% paraformaldehyde (PFA) and embedded in paraffin. Five-μm thick sections were cut and stained with hematoxylin and eosin. Images were acquired using the VS120 slide scanner (Olympus).

### Quantitative RT-PCR of pituitary

Generated knockouts and their siblings were anesthetized, and the pituitary was collected for real-time PCR analysis. Total RNA was extracted from each pituitary using a Fast Gene RNA Basic kit (Nippon Genetics) according to the manufacturer’s protocol. Total RNA was reverse-transcribed with the PrimeScript RT kit (Takara, Kusatsu, Japan) according to the manufacturer’s instructions. For real-time PCR, the cDNA was amplified using KAPA SYBR fast qPCR kit (Nippon genetics) with LightCycler 480 II system (Roche, Mannheim, Germany). The temperature profile of the reaction was 95 °C for 5 min, 45 cycles of denaturation at 95 °C for 10 s, annealing at 60 °C for 10 s, and extension at 72 °C for 10 s. The PCR product was verified by melting curve analysis. A housekeeping gene, β-actin (*actb*) was used for normalization. Primers used in this experiment are shown in Extended Data Table 3.

### Reverse transcription PCR

Adult female d-rR wildtype medaka were deeply anesthetized and their brain and intestine were excised. RNA was extracted using Fast Gene RNA Basic kit according to the manufacturer’s protocol. The RNA was purified with DNase and reverse-transcribed with the PrimeScript RT kit according to the manufacturer’s protocol. The cDNA was amplified using KAPA HiFi Hotstart ReadyMix PCR kit (Nippon Genetics) with T-100 Thermal Cycler (Biorad, Hercules, CA). The temperature profile of the reaction was 95 °C for 2 min, 35 cycles of denaturation at 95 °C for 10 s, annealing at 58 °C for 30 s, and extension at 72 °C for 10 s. The PCR products were verified by gel electrophoresis on 2% agarose gel. The primer pairs used in PCR are listed in Extended Data Table 3.

### Reporter activation assay

The medaka CCK8 (DY(SO_3_H)LGWMDF-NH_2_) and Gastrin8 (DY(SO_3_H)RGWLDF-NH_2_) peptides were synthesized by Scrum (Tokyo, Japan). The cDNA fragment encoding the full-length Cckbr1 was PCR-amplified and subcloned into the expression vector pcDNA3.1/V5-His-TOPO (Thermo Fisher Scientific). The primers used here are listed in Extended Data Table 3. The resulting Cckbr1 expression construct was transiently transfected into Hela cells together with a luciferase reporter vector containing cis-acting elements responsive to cAMP (pGL4.29; Promega), Ca^2+^-dependent nuclear factor of activated T-cells (NFAT) (pGL4.30; Promega), or the MAPK signaling pathway (pGL4.33; Promega) and the internal control vector pGL4.74 (Promega) by using Lipofectamine LTX (Thermo Fisher Scientific). Forty-two hours after transfection, cells were stimulated with CCK8 or Gastrin8 polypeptide at doses of 0, 10^−11^, 10^−10^, 10^−9^, 10^−8^, 10^−7^, and 10^−6^ M for 6 hr. After cell lysis, luciferase activity was measured by using the Dual-Glo Luciferase Assay System (Promega). Each assay was performed in duplicate and repeated three times independently. Hela cells used in this study were authenticated by short tandem repeat profiling (National Institute of Biomedical Innovation, Osaka, Japan) and confirmed to be mycoplasma-free (Biotherapy Institute of Japan, Tokyo, Japan).

### Ca^2+^ imaging

Adult *fsh*:IP and *lh*:IP were anesthetized in MS-222 and were decapitated. The pituitary was excised out to the recording chamber. Unless otherwise mentioned, female fish were used in this experiment. Their body weight was 0.152-0.224g. The medaka CCK8 peptide was diluted to 0.1, 1, 10, 100, or 1000 nM with 0.01% DMSO in ACSF. The pituitary was perfused using a peristaltic pump (Rainin Dynamax RP-1, Rainin, Columbus, OH), and images were taken every 5 seconds with a scientific complementary metal oxide semiconductor camera (scMOS) (Andor Zyla 4.2 PLUS, Oxford Instruments, Belfast, UK) configured to the fluorescent lamp (X-Cite 110 LED Illumination system, Excelitas Technologies, Waltham, MA). Images were captured by Micro-Manager 1.4. The pituitary was washed with ACSF for 10 minutes after CCK8 is applied.

### Incubation of the pituitary followed by quantification of gonadotropin mRNA

Adult female d-rR wildtype and generated *cckbr1* KO medaka were deeply anesthetized and their pituitary was incubated. Their body weight was 0.17-0.36g. We used 0, 10, or 100 nM medaka CCK8 to 200 uL culture medium Leibovitz L-15 medium (Thermo Fisher) supplemented with 1% 100x penicillin-streptomycin, 5% heat-inactivated fetal bovine serum (FBS), and 10 mM D-glucose. The pituitary was incubated at 27 °C for 48 hours. After 48 hours, the medium was removed and qRT-PCR was conducted previously mentioned above.

### Data analysis

All values are shown as mean ± standard error of the mean (SEM). For statistical analyses, groups of two were processed with Student’s *t*-test, and groups of more than three were processed with Dunnett’s test.

Microphotographs and images were processed by ImageJ software (National Institutes of Health, Bethesda, MD). Statistical analyses were performed with Kyplot 6.0 (Kyence, Tokyo, Japan). Graphs were drawn with Kyplot 6.0 or R (R Foundation).

## Acknowledgments

We thank Dr. Shin-ichi Higashijima (National Institute for Basic Biology, Japan) for providing a plasmid containing the medaka heat shock promoter sequence. We also thank the staff at Laboratory for Phyloinformatics, RIKEN BDR for helpful advice on low-input RNA-seq technique. We are grateful to Dr. Daichi Kayo (Tohoku University, Japan) for the helpful discussion and comments. We also thank Dr. Soma Tomihara (Nagahama Institute of Bio-Science and Technology, Japan) for help in the construction of plasmid DNA. This work was funded by the Japan Society for the Promotion of Science for SKa and KO (23H02306) and for SKa (18K19323, 18H04881), and by Mitsubishi Foundation and Sumitomo Foundation for SKa.

## Author contributions

SKU, YN, KO, and SKa conceived the project. SKU, YN, and SKa performed histological analyses, while SKU performed Ca^2+^ imaging and *in vitro* analysis of the pituitary. YN analyzed the knockouts and reporter assay. SKa generated GFP transgenic medaka. KM performed FSH cell–specific RNA-seq. TK and SKu helped with the methodology of FSH cell–specific RNA-seq. KO and SKa assisted with data interpretation. SKU, YN, and SKa wrote the original draft. All authors contributed to the editing of the manuscript.

## Data and materials availability

All data are available in the main text or in the extended data.

