## Supplemental Materials for "Identification of the FSH-RH, the other gonadotropin-releasing hormone"

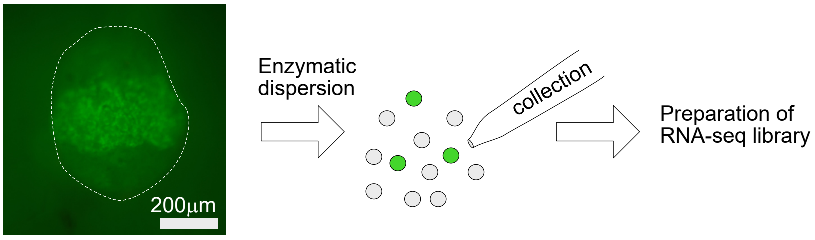


**Extended Data Fig. 1. Schematic of FSH-specific RNA-seq by using FSH-GFP transgenic medaka.**
Left fluorescent image is the pituitary of the FSH-GFP transgenic medaka. Sub-region where FSH cells are localized show GFP fluorescence. Dotted line indicates the edge of the whole pituitary. The pituitary of the FSH-GFP transgenic medaka was excised and subjected to enzymatic dispersion. Then, the GFP positive FSH cells were collected and RNA-seq was performed.

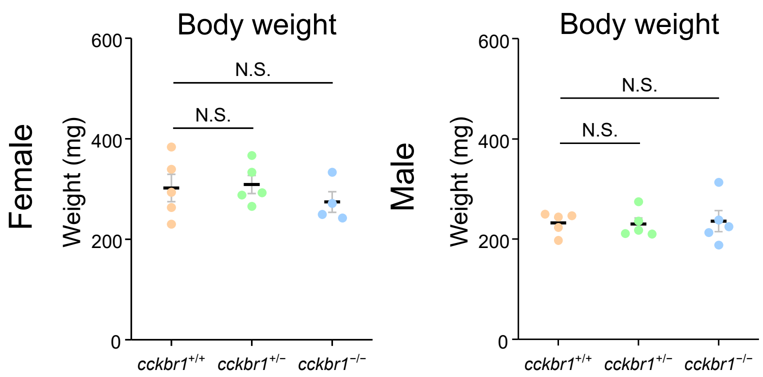


**Extended Data Fig. 2. The body weight does not change among *cckbr1*$^{+/+}$, *cckbr1*$^{+/-}$, and *cckbr1*$^{-/-}$.** Body weight of *cckbr1*$^{+/+}$, *cckbr1*$^{+/-}$, and *cckbr1*$^{-/-}$ female and male medaka. No significant change of body weight was observed among them. N.S., Not significant.

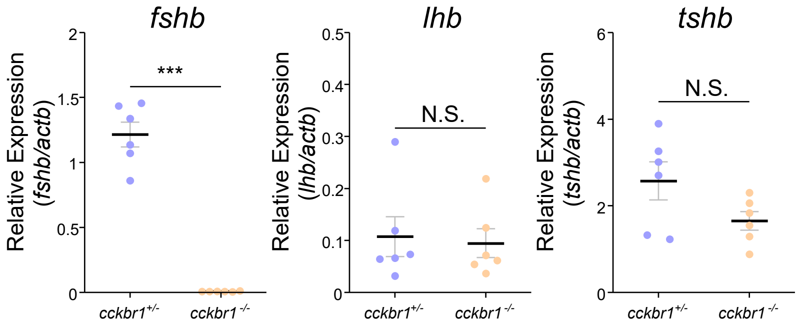


**Extended Data Fig. 3. *fshb* expression decreased in ovarectomized *cckbr1*$^{-/-}$ female.**
The effect of *cckbr1* knockout in ovariectomized (OVX) females. *cckbr1*$^{-/-}$ medaka showed drastic decrease in *fshb*, which is similar to the results of intact females. On the other hand, the effect of *cckbr1* was not observed in *lhb* expression in OVX females. There was no significant difference in *tshb* expression.

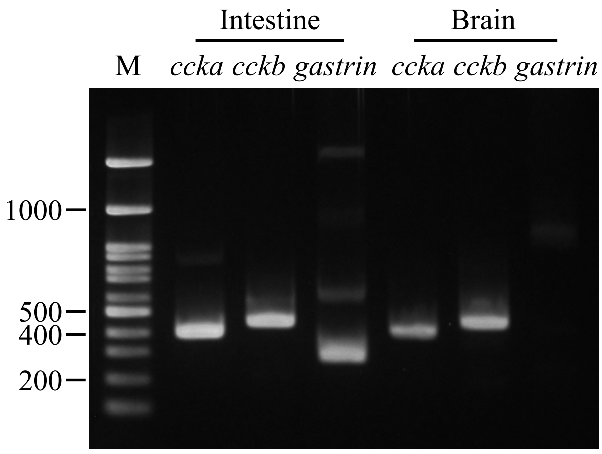


**Extended Data Fig.4.** ***ccka*** **and** ***cckb*** **but not** ***gastrin*** **is expressed in the brain.**
Reverse transcription polymerase chain reaction (RT-PCR) analysis of the medaka brain and intestine. cDNA of brain and intestine were examined for PCR using *ccka*- (381-bp), *cckb*- (437-bp), and *gastrin*- (295-bp) specific primers and resolved in 2% agarose gel electrophoresis. *ccka* and *cckb* were amplified from both brain and intestine cDNA, whereas *gastrin* was amplified only from the intestine cDNA .

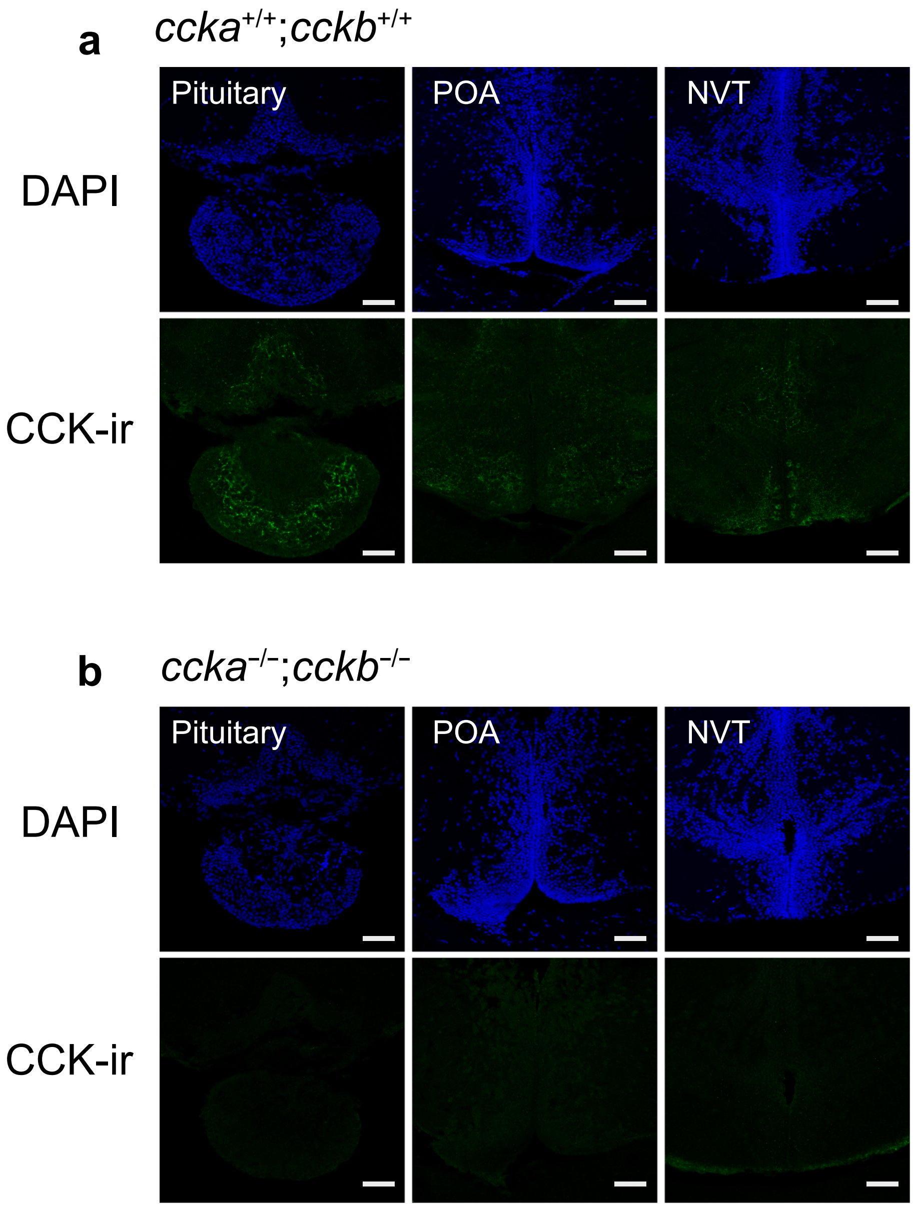


**Extended Data Fig. 5. Cell body and fibers are not observed when knocking out *ccka* and *cckb*.**
**a**, Immunohistochemistry of *ccka* and *cckb* double knockout medaka, using CCK antibody. Cell body and fibers were observed in the pituitary, preoptic area (POA), and nucleus ventralis tuberis (NVT) of *ccka*$^{+/+}$;*cckb*$^{+/+}$ medaka. **b**, On the other hand, in *ccka*$^{-/-}$;*cckb*$^{-/-}$ medaka, no cell body or fibers were observed. Scale bars, 50mm.

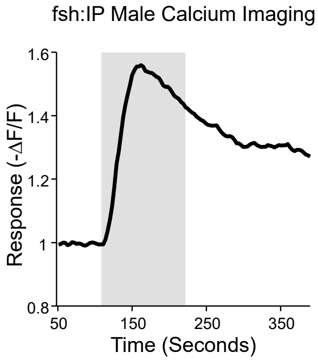


**Extended Data Fig. 6. Increased $[Ca^{2+}]_i$ in FSH cells in male in response to CCK8 peptide.**

FSH cells in males also show $Ca^{2+}$ response to 1000 nM CCK8 peptide. During the application of peptide, FSH cells showed drastic increase of $[Ca^{2+}]_i$ which is similar to the response in females.

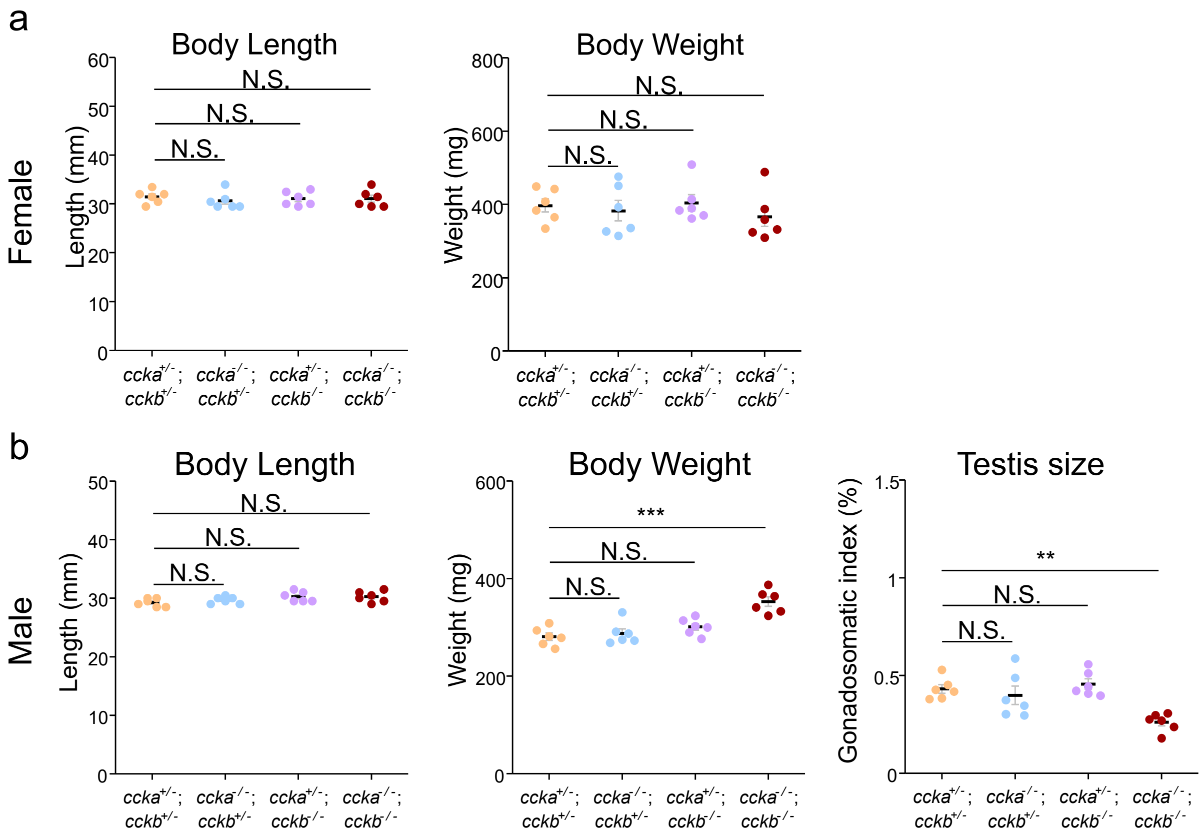


**Extended Data Fig7. Body length and weight is mostly non-significant among *ccka* and *cckb* double KO.**

**a**, The body length and weight of *ccka*$^{+/-}$;*cckb*$^{+/-}$, *ccka*$^{-/-}$;*cckb*$^{+/-}$, *ccka*$^{+/-}$;*cckb*$^{-/-}$, and *ccka*$^{-/-}$;*cckb*$^{-/-}$ female medaka. No significant difference is seen among the KO. **b**, The body length, weight, and gonadosomatic index (GSI) of male *ccka*/*cckb* double KO. The GSI of *ccka*$^{-/-}$;*cckb*$^{-/-}$ was lower among other KO similar to female *ccka*$^{-/-}$;*cckb*$^{-/-}$. ***$P$<0.001, **$P$<0.01, N.S., Not significant.

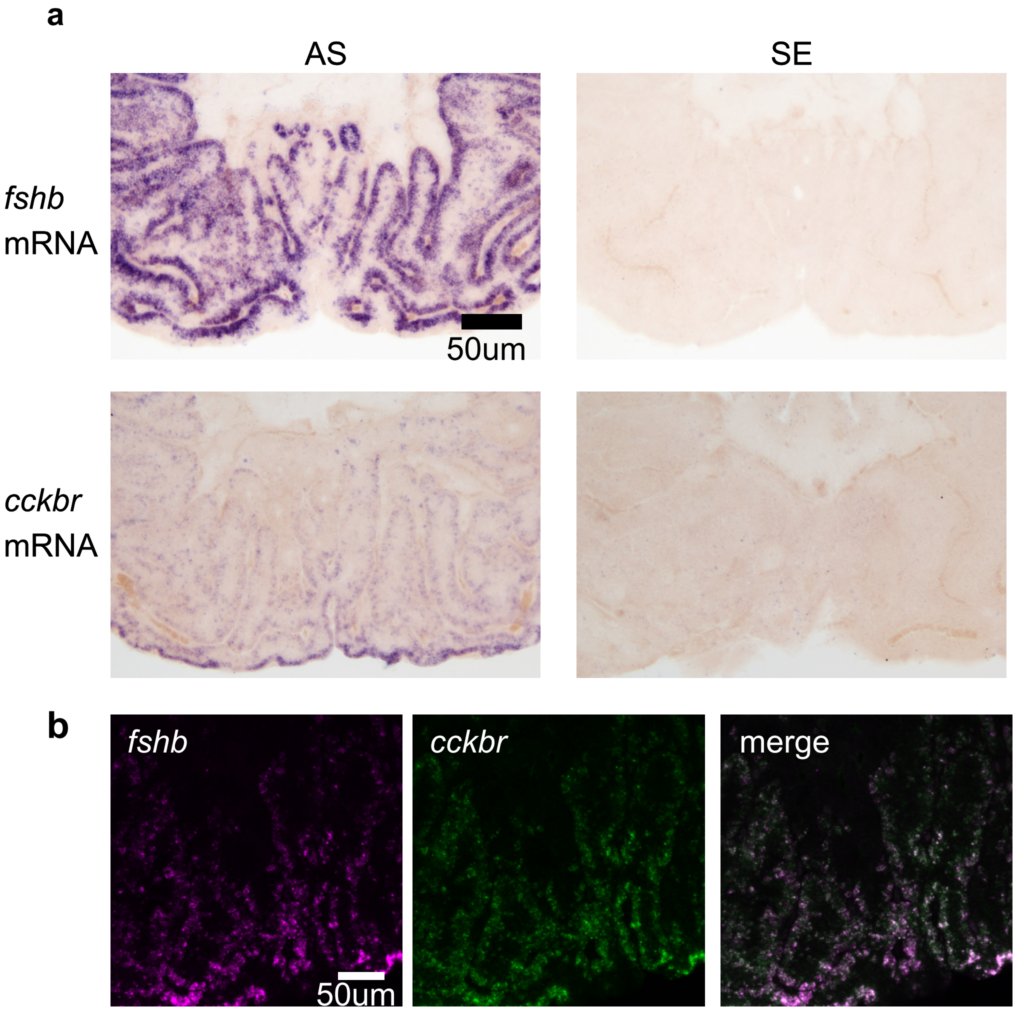


**Extended Data Fig. 8. *fshb* and *cckbr* is expressed in the pituitary of Japanese eel.** In Japanese eel, FSH cells express *cckbr*. **a**, Analysis of expression of *fshb* and *cckbr* by *in situ* hybridization in the pituitary. Only sections hybridized with anti-sense probe were labeled for both *fshb* and *cckbr*. **b**, Co-expression of *fshb and cckbr* in the pituitary by double *in situ* hybridization.

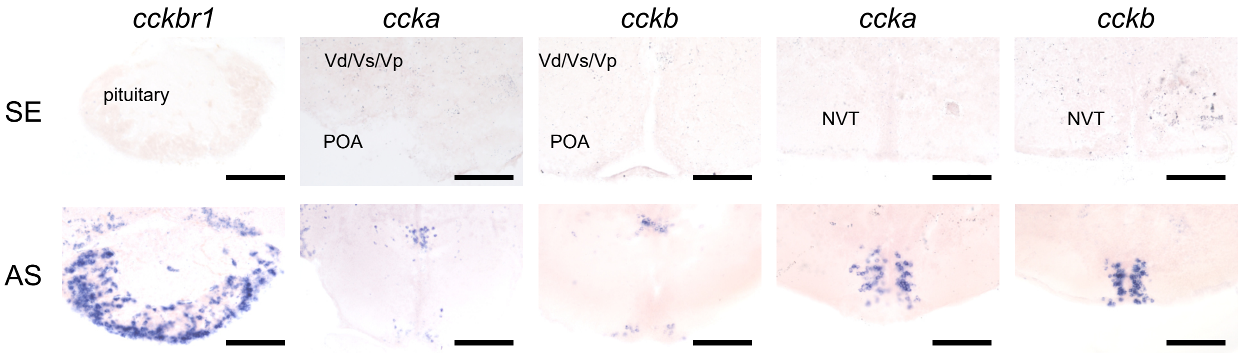


**Extended Data Fig. 9. Specific *in situ* hybridization of *cckbr1*, *ccka*, and *cckb*.**
*in situ* hybridization of *cckbr1*, *ccka*, and *cckb*. Only the sections hybridized with anti-sense probes (AS) were labeled whereas sections hybridized with sense probes (SE) did not show any signal which supports the validity of this labeling. Scale bars, 100 μm.

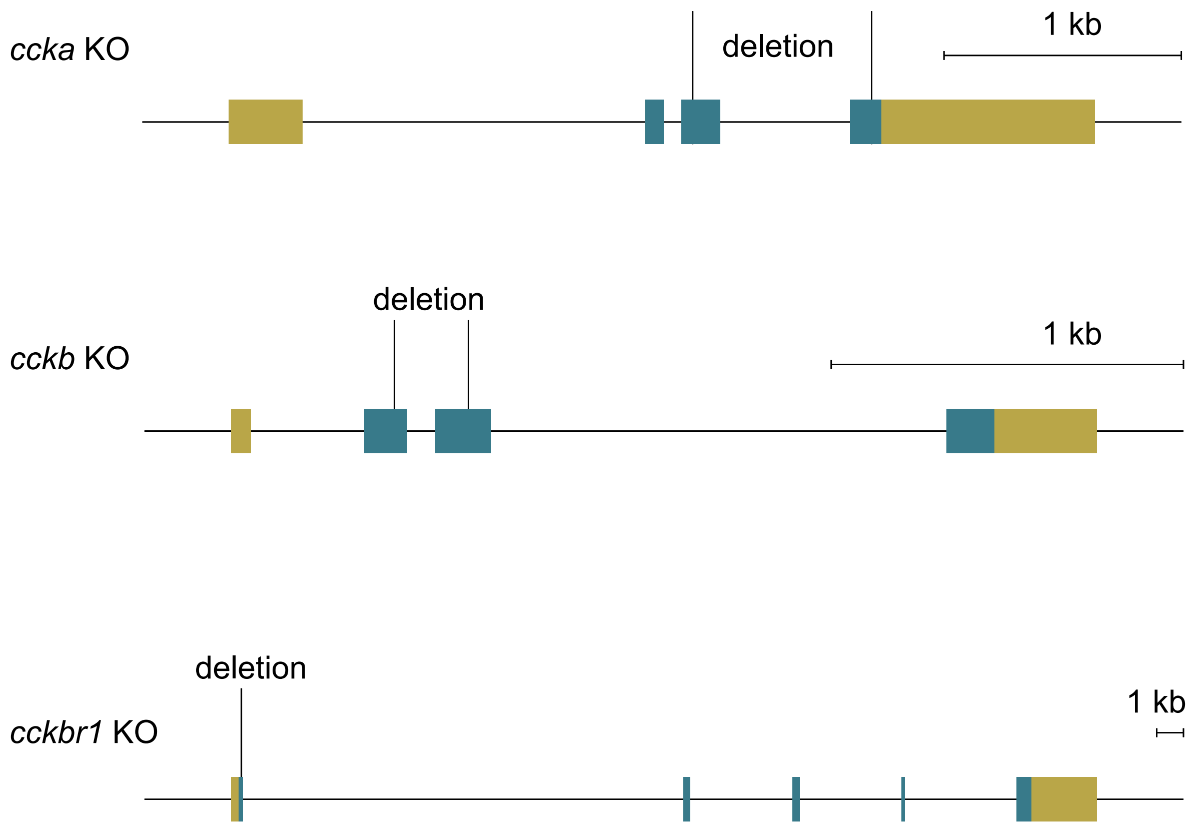


**Extended Data Fig. 10. Knockout lines of medaka.**
Knockout medaka generated in this study. *ccka, cckb,* and *cckbr1* knockout has a 754-bp, 210-bp, and 8-bp deletion, respectively in the area shown in this figure. Note that these deletions induced frame shift in their mRNA sequence.

Extended Data Table1.

| gene_id | Gene name | annotation | Median TPM value of FSH cell pools (n=4) |
|---|---|---|---|
| ENSORLG00000017966 | cckbr1 | cholecystokinin receptor-like | 542.79 |
| ENSORLG00000019757 | gnrh-r2 | gonadotropin-releasing hormone receptor 2 | 449.315 |
| ENSORLG00000016288 | tgfbr3 | transforming growth factor beta receptor type 3 | 442.8 |
| ENSORLG00000000865 | chrna4b | neuronal acetylcholine receptor subunit alpha-2 | 267.245 |
| ENSORLG00000014011 | pgrmc1 | progesterone receptor membrane component 1 | 247.1 |
| ENSORLG00000030131 | pk1-r | pyrokinin-1 receptor-like | 224.915 |
| ENSORLG00000024060 | nr1d2b | nuclear receptor subfamily 1 group D member 2 | 199.77 |

List of highly expressed receptors in the RNAseq analysis of FSH cells

Extended Data Table 2. EC50 values of the reporter assay using Hela cells expressing Cckbr1.

| EC50 | CCK8 | Gastrin8 |
|---|---|---|
| pGL4.29(cAMP) | 0.2 nM | 9.1 nM |
| pGL4.30($Ca^{2+}$) | 0.7 nM | 1.8 nM |
| pGL4.33(MAPK) | 1.0 nM | 1.8 nM |

Extended Data Table 3. Primers used in this study.

| target | direction | purpose | sequence (5' to 3') |
|---|---|---|---|
| *cckbr1* | forward | genotyping (gDNA PCR) | AAAGACGGAGAGCCAAAGACAG |
| *cckbr1* | reverse | genotyping (gDNA PCR) | TGCAGAGATAGCTTTCTCCAAGAT |
| *cckbr1* | forward | genotyping (HRM) | CTGATGGAGCAGCTTCAGAGC |
| *cckbr1* | reverse | genotyping (HRM) | TGCTGCGCTTCTCCTGTCGT |
| *cckbr1* | forward | genotyping (HRM) | ATCTCCTGCGTGAACCCGTCCACGCTT |
| *ccka* | forward | genotyping (gDNA PCR) | GTGTCTAATCCCAATTCAGAAAGTG |
| *ccka* | reverse | genotyping (gDNA PCR) | AAGCTGACAAGCTGTGCCTCTT |
| *ccka* | forward | genotyping (CS) | ATCCTGGCTGCTCTTTTCACTG |
| *cckb* | forward | genotyping (gDNA PCR) | AGCTCCTCACATGACTGAAGCT |
| *cckb* | reverse | genotyping (gDNA PCR) | AGTGGCAGGAAAAAGCACTCG |
| *cckb* | forward | genotyping (CS) | AAACGCTCCGTCTTCTGTCTGTG |
| *cckbr1* | forward | Probe template | CTGATCTCCAGGGAACTTTATCG |
| *cckbr1* | reverse | Probe template | GCAGGAGAAAGTATGGAGTACAG |
| *ccka* | forward | Probe template | ACTGTTTGAAAGCCTCAGCACCA |
| *ccka* | reverse | Probe template | CCCTAGTAGATGATTTGATATGAAGATT |
| *cckb* | forward | Probe template | GAACTGCTCTCCTCACTCTCATA |
| *cckb* | reverse | Probe template | GCAGCGAAGCAGCTTTTGCTG |
| *ccka* | forward | RT-PCR | GCAGTCATGAATGTAGGAATCTACGT |
| *ccka* | reverse | RT-PCR | TTATGGGGAATATTCATACTCCTCCACAC |
| *cckb* | forward | RT-PCR | TTCTCTCCTCAAGATGACCGCTG |
| *cckb* | reverse | RT-PCR | AAGAGTACGAGTACTCCTCATAAGGG |
| *gastrin* | forward | RT-PCR | AGGCAGCCATGTCAGGGAAAAC |
| *gastrin* | reverse | RT-PCR | TCCCTCCTCCCAAAGTCCA |
| *eel_fshb* | forward | Probe template for eel | CAGATTCACAGTTGCCATGCATCT |
| *eel_fshb* | reverse | Probe template for eel | CATCTATCCCTTGCCGCAGTT |
| *eel_cckbr* | forward | Probe template for eel | GCGATGGACGTACAGAAACTGAATG |
| *eel_cckbr* | reverse | Probe template for eel | CTTCCAGGTGTTGACGGAGTAG |
| *cckbr1* | forward | cds cloning (including sequence for construction) | CTTGCCGCCATGGATACTTTGAGAAACG AGAC |
| *cckbr1* | reverse | cds cloning (including sequence for construction) | CTCTCAGCAGTTTCCCATGGTG |
| *actb* | forward | qPCR | GTGATGTTGATATCCGTAAGGATCTGTA |
| *actb* | reverse | qPCR | TCTGGTGGGGCAATGATCTTGA |
| *fshb* | forward | qPCR | TGGAGATCTACAGGCGTCGGTAC |

| | | | |
|---|---|---|---|
| *fshb* | reverse | qPCR | AGCTCTCCACAGGGATGCTG |
| *lhb* | forward | qPCR | TGCCTTACCAAGGACCCCTTGATG |
| *lhb* | reverse | qPCR | AGGGTATGTGACTGACGGATCCAC |
| *tshb* | forward | qPCR | GCTACTCAAGGGACAGCA |
| *tshb* | reverse | qPCR | GCAGCCTCTCTGGATAAGGAA |

gDNA PCR, PCR on genomic DNA; CS, cycle sequence; HRM, high-resolution melt analysis. Unless otherwise specified, all target genes mentioned are from the medaka.
